# The Immunomodulatory Effects of Social Isolation in Mice are Linked to Temperature Control

**DOI:** 10.1101/2021.09.03.458884

**Authors:** Alice Hamilton, Raffaella Rizzo, Samuel Brod, Masahiro Ono, Mauro Perretti, Dianne Cooper, Fulvio D’Acquisto

## Abstract

Living in isolation is considered an emerging societal problem that negatively affects the physical wellbeing of its sufferers in ways that we are just starting to appreciate. This study investigates the immunomodulatory effects of social isolation in mice, utilising a two-week program of sole cage occupancy followed by the testing of immune-inflammatory resilience to bacterial sepsis. Our results revealed that mice housed in social isolation showed an increased ability to clear bacterial infection compared to control socially housed animals. These effects were associated with specific changes in whole blood gene expression profile and an increased production of classical pro-inflammatory cytokines. Interestingly, equipping socially isolated mice with artificial nests as a substitute for their natural huddling behaviour reversed the increased resistance to bacterial sepsis. These results further highlight the ability of the immune system to act as a sensor of our living conditions and to respond in a compensatory fashion to external challenges that might threaten the survival of the host.

## Introduction

Increasing evidence is showing the immune system as a sensor of a wide range of changes to a healthy state of homeostasis (1-3). These include classical challenges coming from pathogens, organ disfunction caused by tissue and cell damage, and any other changes that can occur ‘outside’ the physical body. These are both positive and negative conditions like fear (4, 5), anger (6-8), stress (9-12), a state of eudaemonia (13-15), falling in love (16) or general relaxation (17-21). Unlike the classical challenges described above, these external conditions modulate the immune response in ways that are far from being fully established. Indeed, there are no receptors or molecules that specifically respond to changes in external conditions but rather a concerted response of different cells and tissues - for instance, the hypothalamic-pituitary-adrenal (HPA) axis and the sympathetic nervous system (SNS) - that ultimately impact the host immune response at multiple levels (22-25).

Clinical experimental evidence suggests that social connections and a sense of belonging are key factors in determining the severity and development of a wide range of pathologies including immune disorders (26-29). Studies have suggested that a lack of social connections could be as damaging to health as smoking 15 cigarettes per day (30). Investigations in experimental animal models and humans have shown that poor social connections affect the body at the behavioural (31, 32), psychological (33, 34) and physiological levels (35-37). All of which accumulate to increase the risk of chronic inflammatory diseases such as Alzheimer’s disease and cardiovascular disease and all-cause mortality (6–10). In humans, poor social connections result in an increased risk of depression (38-40) and anxiety which can lead to a cascade of poor lifestyle choices including a sedentary lifestyle, increased drinking, smoking and poor diet (41-45) and cause further withdrawal from society (11). This is clearly a vicious cycle.

We are interested in understanding how external living conditions, lifestyle and emotions impact the immune response and specifically if each of these conditions influences (or not) the immune response in a unique way (9, 10, 46-48). In previous studies carried out by our group (49), it was shown that just 2 weeks of environmental enrichment in mice was sufficient to alter the immune cell response to inflammatory stimuli both in the *in vitro* and *ex vivo* settings. At the basal level, it was demonstrated that environmental enrichment increased the overall blood cellularity in comparison to control mice. When challenged with zymosan-induced peritonitis, environmentally enriched mice had enhanced neutrophil and macrophage recruitment to the peritoneum compared to socially housed mice. Using a model of caecal ligation and puncture it was demonstrated that environmentally enriched mice showed enhanced systemic microbial clearance compared to standard housed mice. Interestingly, this improved host response was linked to increased cellularity of the immune system and to a specific pattern of gene expression of the whole blood that led to an enhanced and immune-protective immune response to infection.

In this study, instead of giving mice socially and physically stimulating housing conditions such as that used in environmental enrichment, we asked what would happen to the immune-inflammatory response if a polar opposite housing condition was used. Unexpectedly, the results showed that similarly to the enriched environment, social isolation primes the immune system ready for infection allowing socially isolated (SI) mice to clear bacterial challenges much more effectively than socially housed (SH) mice. Unlike enriched mice, however, the increased host response of SI mice was associated with elevated production of inflammatory cytokines. In addition to this, our data suggest that the loss of social thermoregulation might account for the immunological changes exhibited during social isolation as these changes were partially reversed when the SI mice were given the option of using an artificial nest to keep themselves warm. Overall, our findings provide further evidence of the immune system as a “mirror” and a sensor of changes in living conditions (10).

## RESULTS

### Weight Gain and Food Intake in Socially Isolated Mice

We began our study by assessing the effects of sole living on the general welfare of CD-1 mice. Monitoring weight gain and food/water intake over the 2-week period we observed that SI mice consistently gained ∼30% less weight (SH: 6.45±0.28g *vs* SI: 4.99±0.17g; p<0.0002) compared to SH mice (**Figure 1A**). To determine whether the difference in weight gain was a consequence of reduced food and water intake, we measured the average food and water consumption per mouse in SH and SI animals on day 7 and day 14. SI mice consumed about ∼37% more food compared to SH mice (SH: 70.9±1.78g *vs* SI: 97.5±3.09g; p<0.0001) (**Figure 1B**, left panel). Water consumption between the housing groups (SH: 102±3.04g *vs* SI: 99.7±3.88g) was similar (**Figure 1B**, right panel).

**Figure 1.**
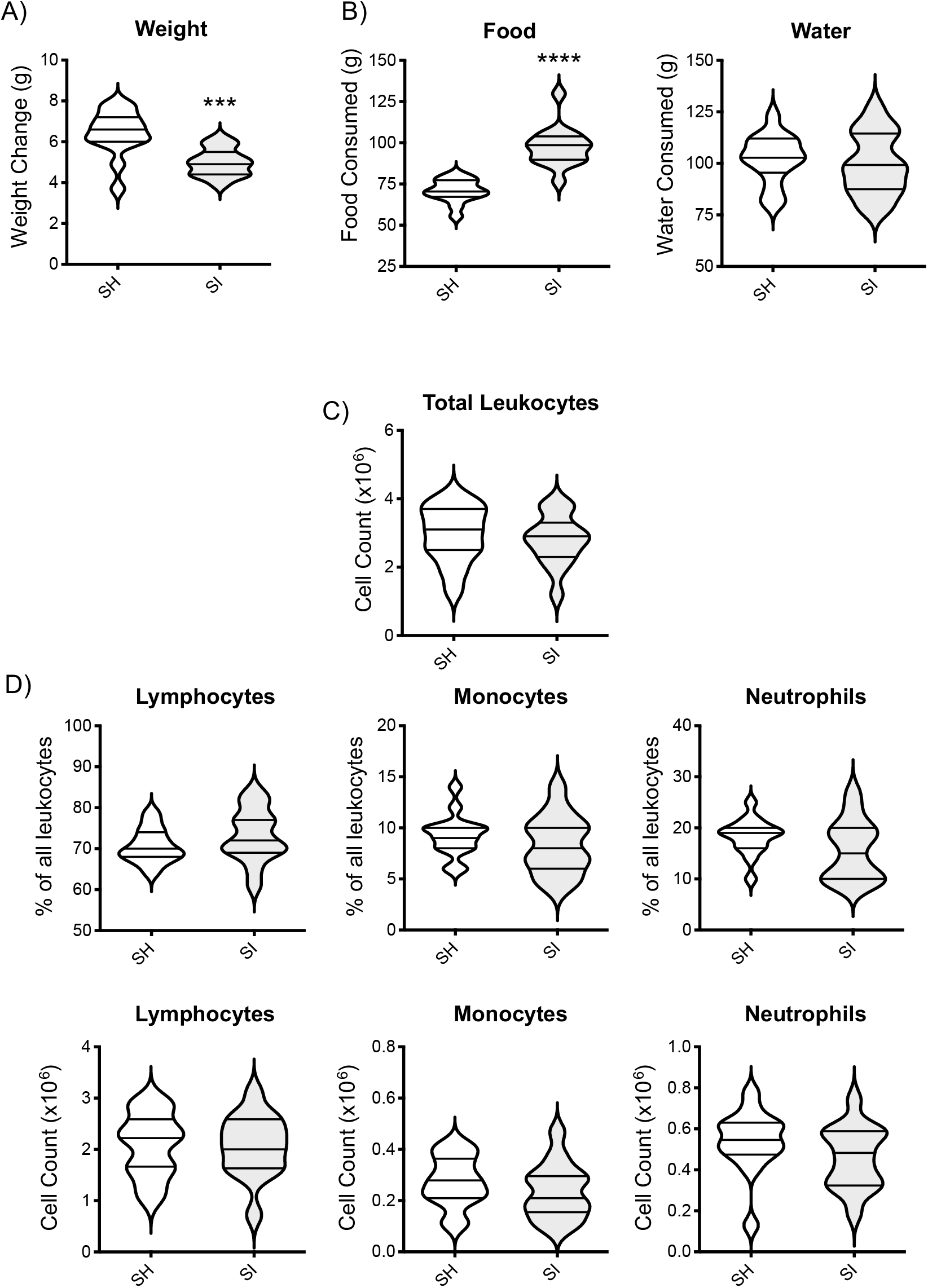
Effect of social isolation on metabolic and blood parameters in CD-1 mice. The violin plots show the net weight gain (A) and food and water intake (B) of CD-1 mice after 2 weeks of social isolation (SI) or social housing (SH). The violin plot in (C) shows the total number of circulating leukocytes from the same mice while the plots in (D) show the relative % or the total cell number of the 3 main blood leukocyte populations. Each plot shows the median and the quartile of n=15 mice. Data are representative of n=3 independent experiments with similar results. ***p<0.001; ****p<0.0001 (Student’s t-test) indicate significant values compared to socially housed mice.

We wondered if food consumption might be linked to any change in stress levels and hence, we conducted standard behavioural tests for anxiety-like behaviour: namely the open field tests (50) and the light and dark box test (51, 52). In agreement with other studies (53, 54), we found significant differences between SH and SI mice in the open field test including an increase in overall horizontal activity (evaluated by the number of squares crossed) and a reduced number of centre crossing in the latter compared to the former (**Supplementary Figure 1A**). Similarly, in the light/dark shuttle box, the number of crossing or time in the light (both parameters of anxiety-like-behaviour) were significantly reduced in SI mice compared to SH control (**Supplementary Figure 1B**).

### Blood Cellularity and Biochemistry of Socially Isolated Mice

We next assessed the impact of social isolation on the overall immune repertoire of CD-1 mice. SI mice showed no significant differences in the overall number of blood circulating leukocytes compared to SH mice (SH: 2.95×10^6^±0.21 *vs* SI: 2.81×10^6^±0.18 cells/ml) (**Figure 1C)**. Similarly, there were no significant differences in the percentages (top panels) or total number (bottom panels) of lymphocytes, monocytes or neutrophils between the two housing groups (**Figure 1D**).

To further ascertain the absence of any type of sickness or Impaired welfare we measured circulating levels of alanine transaminase (ALT), aspartate transaminase (AST), creatinine, and glucose as biomarkers of organ damage or general inflammation. No significant differences were observed in plasma ALT and creatinine concentrations or the AST: ALT ratio between the housing groups (**Figure 2A**). An increased concentration (∼67%) of plasma glucose was seen in SI mice compared to SH mice (SH: 9.20±0.38mmol/L *vs* SI: 15.4±0.91mmol/L; p<0.0001) (**Figure 2B**, middle panel). Plasma corticosterone, one of the main biomarkers of acute stress (55, 56), was comparable between the two housing groups and no statistical difference was observed (**Figure 2B**, right panel), because of habituation (56).

**Figure 2.**
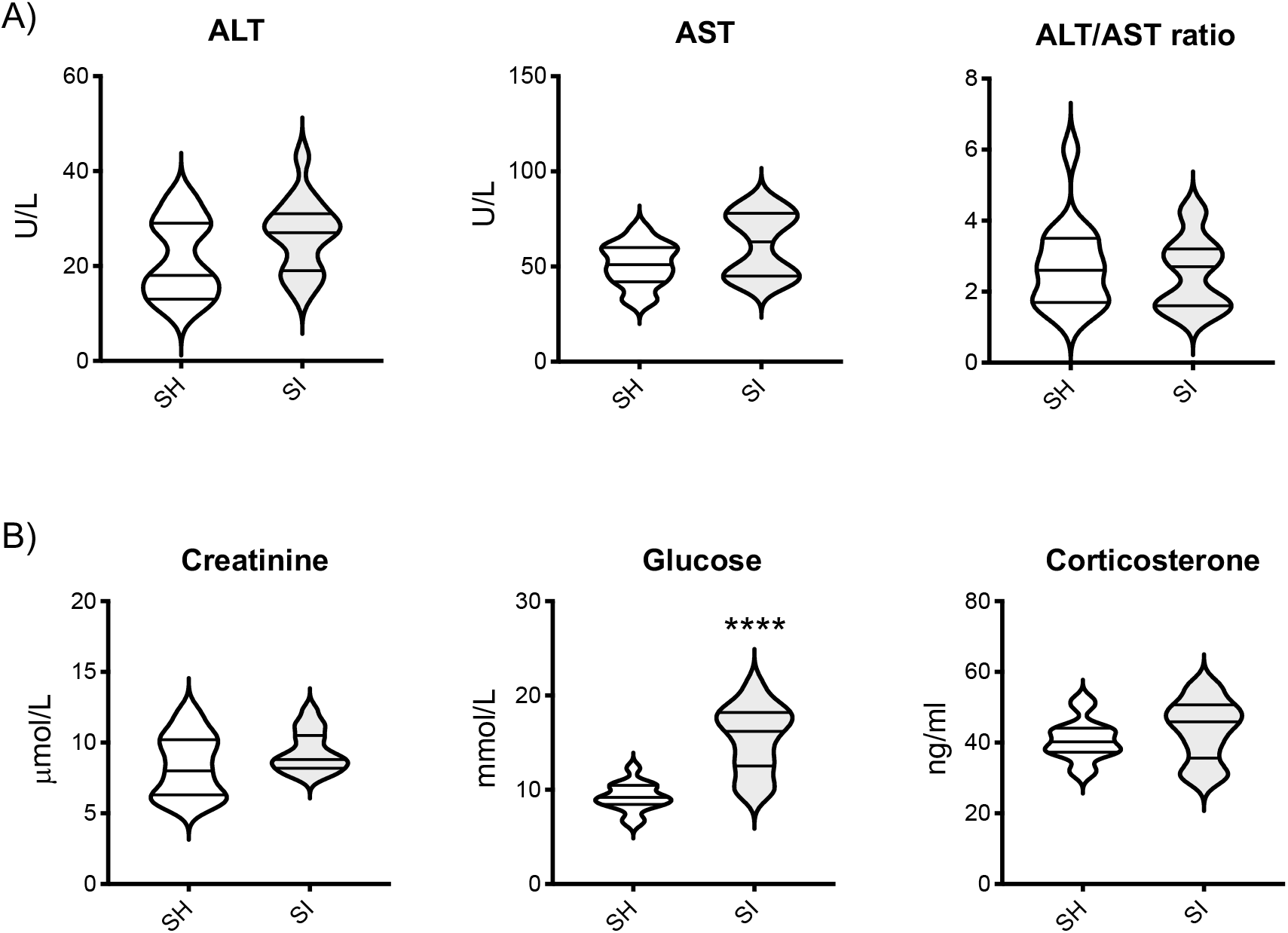
Effect of Social Isolation on the basal blood biochemistry of CD-1 mice. The violin plots in (A) show the levels of ALT, AST, and their ratio in the blood of CD-1 mice after 2 weeks of social isolation (SI) or social housing (SH). The violin plots in (B) show the blood levels of creatinine, glucose or corticosterone in the same mice. Each plot shows the median and the quartile of n=15 mice. Data are representative of n=3 independent experiments with similar results. ****p<0.0001 (Student’s t-test) indicates significant values compared to socially housed control mice.

### Response to Sepsis in Socially Isolated Mice

We explored the response of SI mice to bacterial infection using two similar and well-established experimental stimuli: LPS and *E*.*coli 06:K2:H1* [ATCC®19138™], a live strain of bacteria. In the first series of experiments, mice were culled at 2hr post-sepsis induction as this is the time where the three main inflammatory cytokines we have measured – IL-6, MCP-1 and TNF-α – are known to peak in plasma (57-59). We also measured the same cytokines in the peritoneal lavage fluid (PLF) as a reflection of the inflammatory response at the site of infection. As shown in **Figure 3A**, LPS-challenged SI mice had a higher inflammatory response compared to SH mice as in both peritoneal fluid (bottom panels) and plasma (top panels) there was a significant increase in IL-6, MCP-1 and TNF-α that ranged from about 40 to 80%. Similar findings were observed in SI mice injected with live *E. Coli*: plasma and peritoneal fluids contained significantly higher levels of IL-6, MCP-1 and TNF-α compared to SH mice (**Figure 3B**, top and bottom panels, respectively).

**Figure 3.**
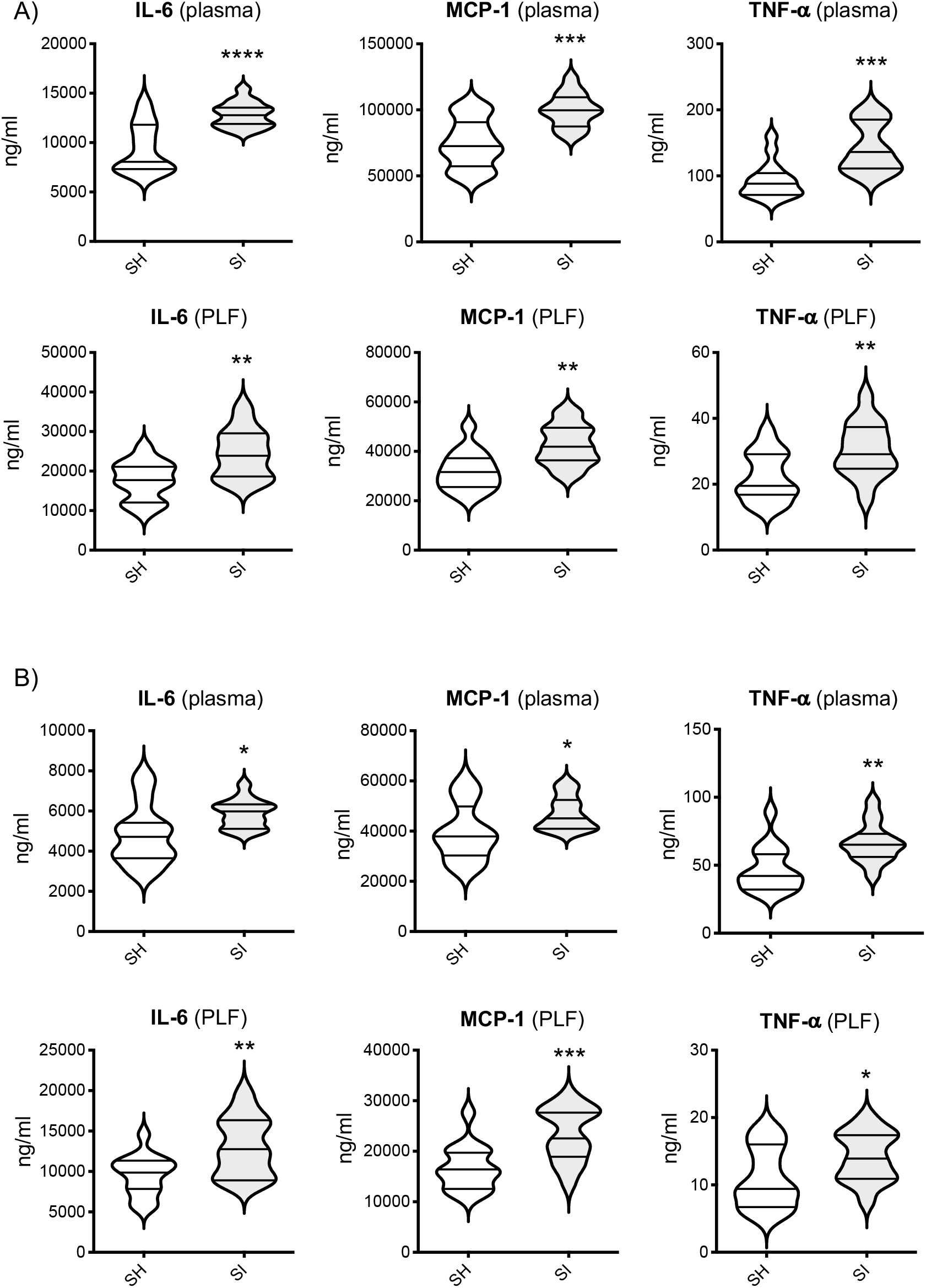
Effect of Social Isolation on LPS or *E. Coli*-induced Inflammation in CD-1 mice. The violin plots in (A) show the levels of IL-6, MCP-1 and TNF-α in the plasma or peritoneal lavage fluid (PLF) of socially isolated (SI) or socially housed (SH) CD-1 mice challenged with LPS (15mg/kg). The violin plots in (B) show the levels of the same cytokines in mice challenged with 1×10^7^ cfu of *E*.*coli 06:K2:H1*. Each plot shows the median and the quartile of n=15 mice. Data are representative of n=3 independent experiments with similar results. *p<0.05; **p<0.01; ***p<0.001; ****p<0.0001 (Student’s t-test) indicates significant values compared to socially housed mice.

### Social Isolation Primes Immune Cells Towards Bacterial Clearance

To investigate the functional implications of the heightened inflammatory response of SI mice, we next assessed bacterial clearance and spreading by measuring bacterial load in plasma and peritoneal fluid at 6 hours post-injection. As shown in **Videos 1 (SH)** and **2 (SI)**, SH mice exhibited the expected systemic clinical signs of sickness such as piloerection, reduced motor activity and lethargy (60-62). Conversely, SI mice exhibited fewer signs of sickness and their motor activity show no impairment. Weight loss over the 6 hours was assessed as an objective measure of sickness (63) and SI mice on average lost significantly less weight compared to SH mice (**Figure 4A**; SH: 1.15±0.11g *vs* SI: 0.69±0.10g; p<0.0052). Bacterial colony counts in blood and PLF of SI mice showed a drastic reduction (p<0.0001) compared to SH animals thus further confirming an enhanced ability of the former to kill pathogens at the site of infection and limit the spreading of infection at the systemic level (**Figure 4B**). Consistent with these data, plasma levels of ALT, AST and creatinine were significantly higher in SH mice compared to SI thus further confirming the presence of infection-driven damage of liver (AST and ALT) and kidney (creatinine) in the former but not the latter (**Figure 4C**).

**Figure 4.**
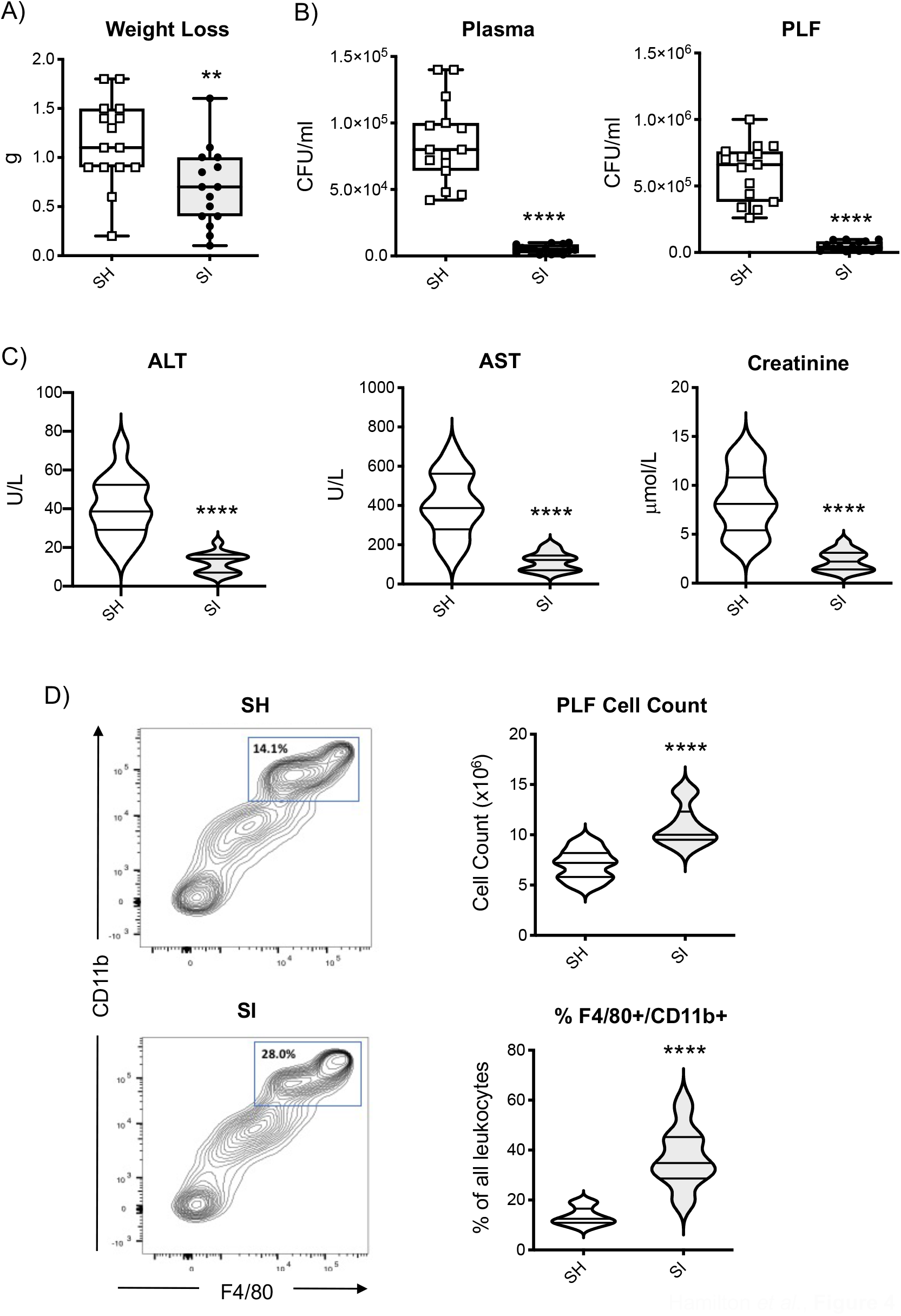
Effect of social isolation on bacterial clearance in CD-1 mice. The box and whisker plots in (A) show the weight loss of socially isolated (SI) or socially housed (SH) CD-1 mice 6 hrs after 1×10^7^ cfu of *E*.*coli 06:K2:H1*challenge. The box and whisker plots in (B) show the bacterial load of blood and peritoneal lavage fluid (PLF) of the same mice while the violin plots in (C) show the blood levels of ALT, AST, and creatinine. The contour plots in (D) show typical staining for CD-11b and F4/80 of peritoneal cells recovered from *E. Coli* challenged mice while the violin plots show the total number of peritoneal cells and their % of gated CD-11b/F4/80^high^ cells. Each plot shows the median and the quartile of n=15 mice. Data are representative of n=3 independent experiments with similar results. **p<0.01; ****p<0.0001 (Student’s t-test) indicates significant values compared to SH mice.

Finally, analysis of the inflammatory cells accumulated at the site of infection revealed another interesting difference between the two groups. The total number of cells found in the peritoneal cavity of SI mice was significantly higher (∼54%) compared to that of SH mice (SH: 7.06×10^6^±0.38 *vs* SI: 10.9×10^6^±0.51 cells/ml). In addition to this, phenotypic analysis of these cells by flow cytometry revealed that almost double the percentage of these cells expressed high levels of mature monocyte/macrophage markers CD-11b and F4/80, thus provide a possible further explanation for the increased bacterial clearance found in SI mice (**Figure 4D**).

### Social Isolation Induces a Unique Whole Blood Gene Profile

To further explore the effect of social isolation on the whole immune system, we compared the gene expression profile of the whole blood between the two experimental groups using the same microarray analysis we previously used for the enriched environment (49). Fold change (FC) and a non-adjusted p-value of <0.05 was used to identify which of the 34,760 probes present on the chip were found to be upregulated (FC ≥2) or downregulated (FC <0.5). The results showed that SI mice have 26 genes upregulated and 10 genes downregulated in comparison to SH mice (**Figure 5A**). Among the upregulated genes, 8 annotated genes do not have a canonical name (*A630089N07Rik, 1110025L11Rik, 1700097N02Rik, 6820431F20Rik, 1700054O19Rik, Gm6445, Gm1966, GM6793*), 3 miRNA (Mir467e, Mir512, Mir5104), 15 known genes including surface receptors (*CD52, CD55*) intracellular signalling molecules (*Dennd5b, Glb1, Laptm4b, Xaf1, Ifi214, Rnu73B*), carrier (*Slc30a4*) or cytoskeletal (*Dnah8*) proteins, and transcription factors/DNA binding proteins (*Zfp729a, Zfp992, Hist1h2bg, Bach2*). The downregulated genes included a lncRNA (*4933432K03Rik*), a predicted gene (*Igkv10-9*), 3 transcriptional regulators (*Hist1H2bc, Hist1h4k, Hist1h2bf*), a surface receptor (Slamf1), and intracellular signalling molecules (*Rab27b, Ctla2b, Rhoj, Gdpd3*). We assessed the validity of these findings by running a real-time PCR for 3 selected genes - focusing on those that were most interesting in terms of immunological functions. As shown in **Figure 5B**, SI mice showed significantly higher levels of *Bach2, CD52* and *CD55* compared to SH thus confirming the microarray data.

**Figure 5.**
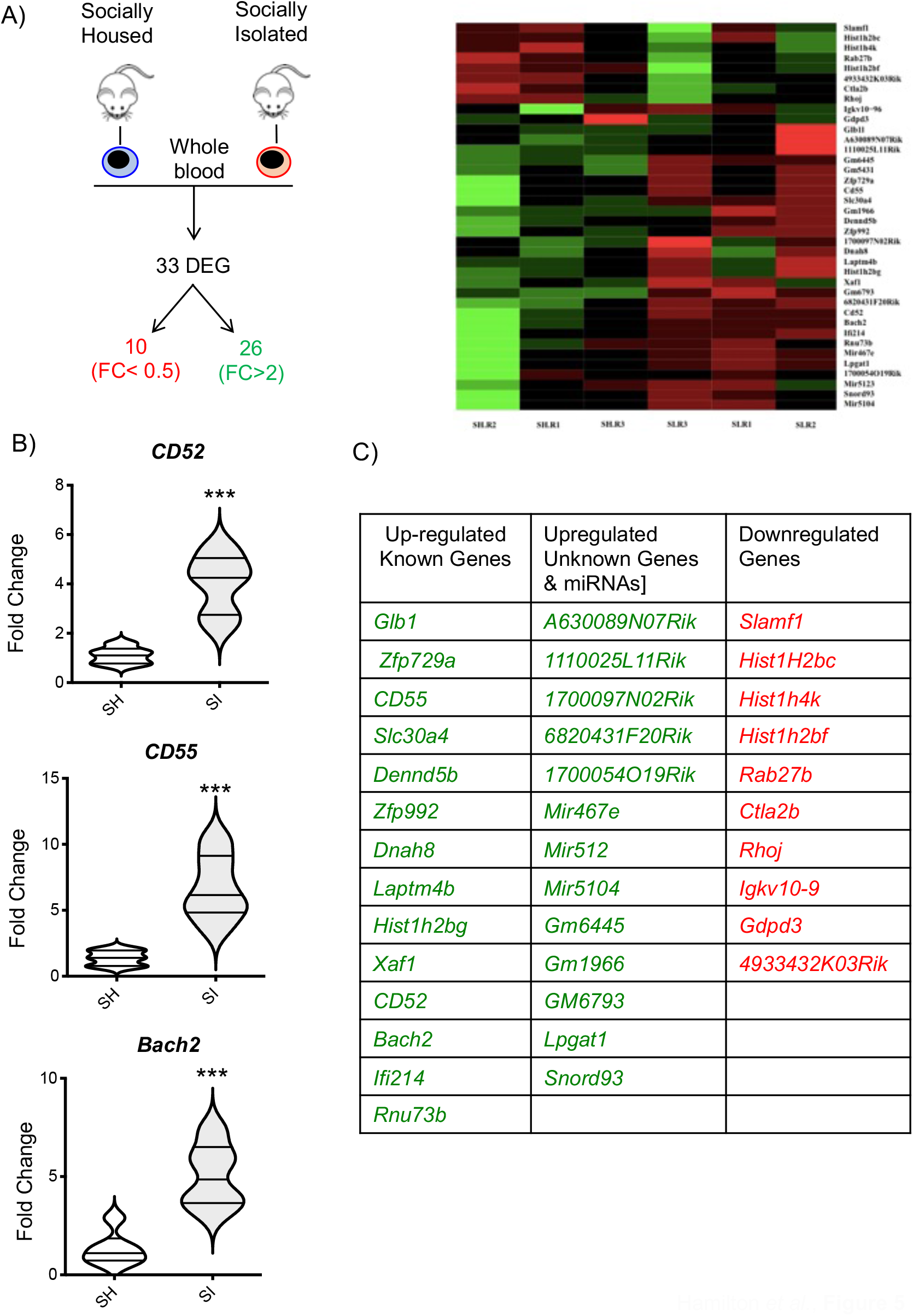
Effect of Social Isolation on the gene expression profile of the whole blood of CD-1 mice. The scheme in (A) provides a graphical representation of the main outcome of the microarray analysis i.e. the total number of differentially regulated genes (DEG) of which 26 were upregulated and 10 downregulated in socially isolated (SI) CD-1 mice compared to control socially housed (SH) mice. The panel on the right is a heatmap analysis of microarray data. The violin plots in (B) show the real-time PCR analysis of 3 genes of interest selected from the microarray analysis. Each plot shows the median and the quartile of n=5 mice. ***p<0.001 (Student’s t-test) indicates significant values compared to socially housed control mice. The table in (C) lists the DEGs obtained from the analysis.

### The Immunomodulatory Effects of Social Isolation are Linked to Thermoregulation

The data shown in **Figure 1** showed that SI mice did not gain extra weight compared to SH despite the significant increase in food consumption. We speculated that this might be due to an increased energy expenditure to keep themselves warm. Indeed, many animals (64-68) including mice (69, 70) use social thermoregulation as a way to maintain a high body temperature without using as much energy as they would on their own. To test this hypothesis, we provided SI mice with an artificial nest that they could use to keep warm at their leisure as they would do in social housing settings. **Figure 6A** shows exemplary photos of the cage set up for SI and SI+Nest mice.

**Figure 6.**
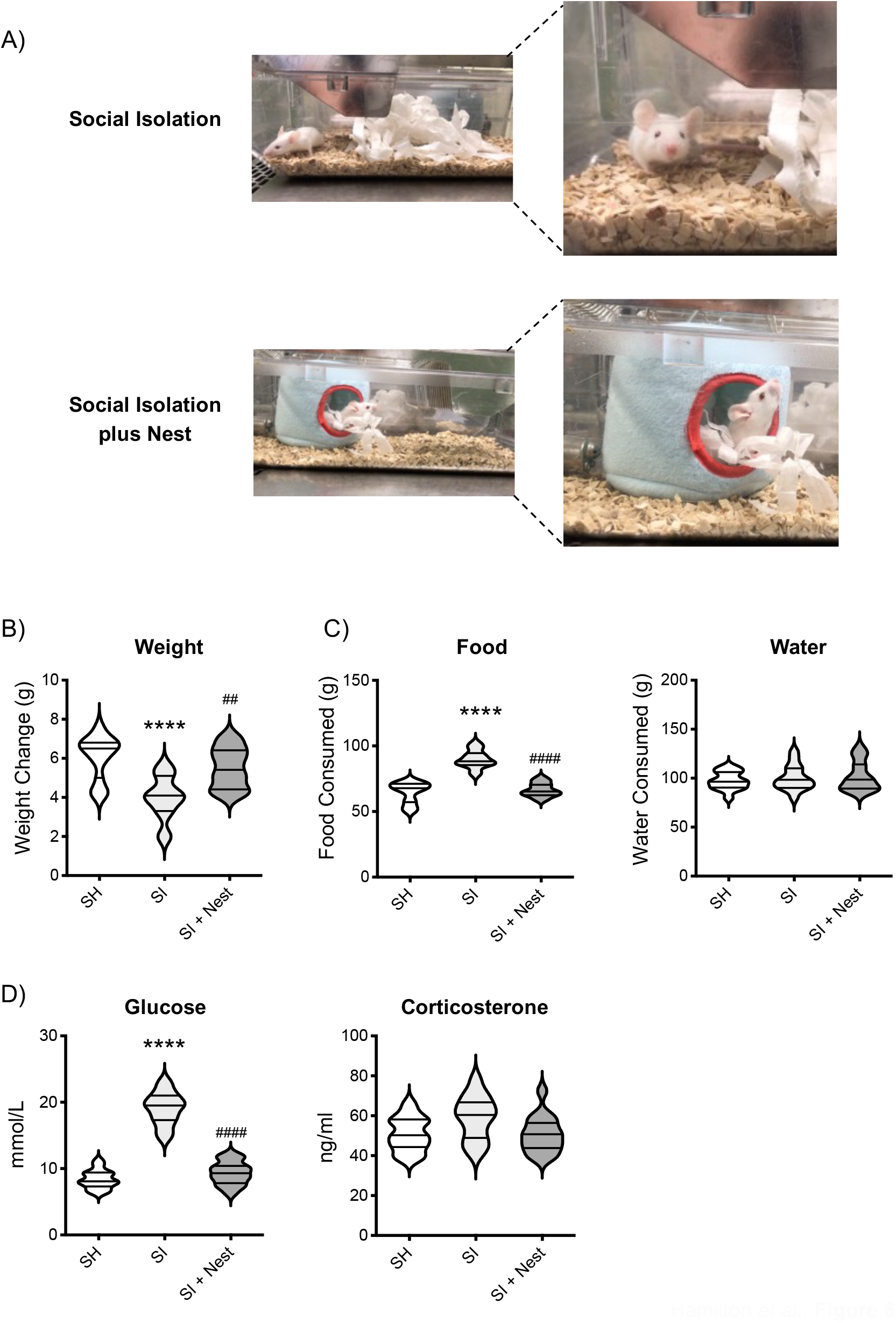
The effects of social isolation on food intake and weight gain in CD-1 mice are reverted by the addition of an artificial nest. The photos in (A) are representative pictures of the cage setting used for social isolation or social isolation + nest. The violin plots show the net weight gain (B) and food and water intake (C) of CD-1 mice after 2 weeks of social isolation (SI), social isolation + nest (SI+Nest) or social housing (SH). The violin plots in (D) show the blood levels of glucose or corticosterone in the same mice. Each plot shows the median and the quartile of n=15 mice. Data are representative of n=3 independent experiments with similar results. ****p<0.0001 (One-way ANOVA) indicates significant values of socially isolated mice compared to socially housed control while ^####^p<0.0001 indicates significant values of SI+Nest mice compared to SI.

First, we assessed whether those mice given an artificial nest had reversed the increase in food consumption. Consistent with our expectation, SI+Nest mice ate significantly less food (p<0.001) than SI mice and consumed a similar amount of food (SH: 64.6±2.04g; SI: 89.8±1.76g; SI+Nest: 66.6±1.34g) to SH mice (**Figure 6C**).

Weight gain reflected these changes and showed similar weight between SH and SI+ Nest mice (SH: 6.07±0.28g; SI: 4.0±0.28g; SI+Nest: 5.5±0.26g) – both been significantly different from SI (**Figure 6B**). Water intake was similar across all three groups (**Figure 6C**). We measured blood glucose levels as this was one of the parameters that were increased by social isolation and this was also reversed by the addition of the artificial nest (**Figure 6D**, left panel; SH: 8.39±0.36mmol/L; SI: 19.1±0.63mmol/L; SI+Nest: 9.33±0.45mmol/L). Conversely, social isolation did not affect plasma levels of corticosterone and this effect was not modified by the addition of the nest (**Figure 6D**, right panel). Similar results were obtained for ALT, AST and creatinine (data not shown).

Next, we addressed the effect of the nest on sepsis and bacterial clearance. The results showed that the weight loss over the 6 hours of *E. Coli* induced sepsis was similar between SH and SI+Nest groups (SH: 1.29±0.08g; SI: 0.65±0.07g; SI+Nest: 1.17±0.06g) and significantly different from that of the SI mice (**Figure 7A**). The analysis of bacterial clearance provided a similar picture with SH and SI+Nest mice showing similar CFU/ml count in both blood and PFL while being significantly higher than in SI animals. (**Figure 7B**). Mirroring these differences, the plasma concentration of ALT, AST and creatinine in SI+Nest mice were also brought back to the same level of SH following the addition of the nest (**Figure 7C**). Analysis of the inflammatory leukocytes accumulated in the peritoneal cavity followed the trend observed for all the other measurements and showed an almost complete reversion of the increase in total cell number and percentages of CD-11b/F4/80^high^ cells in SI mice compared to SI+Nest (**Figure 7D**).

**Figure 7.**
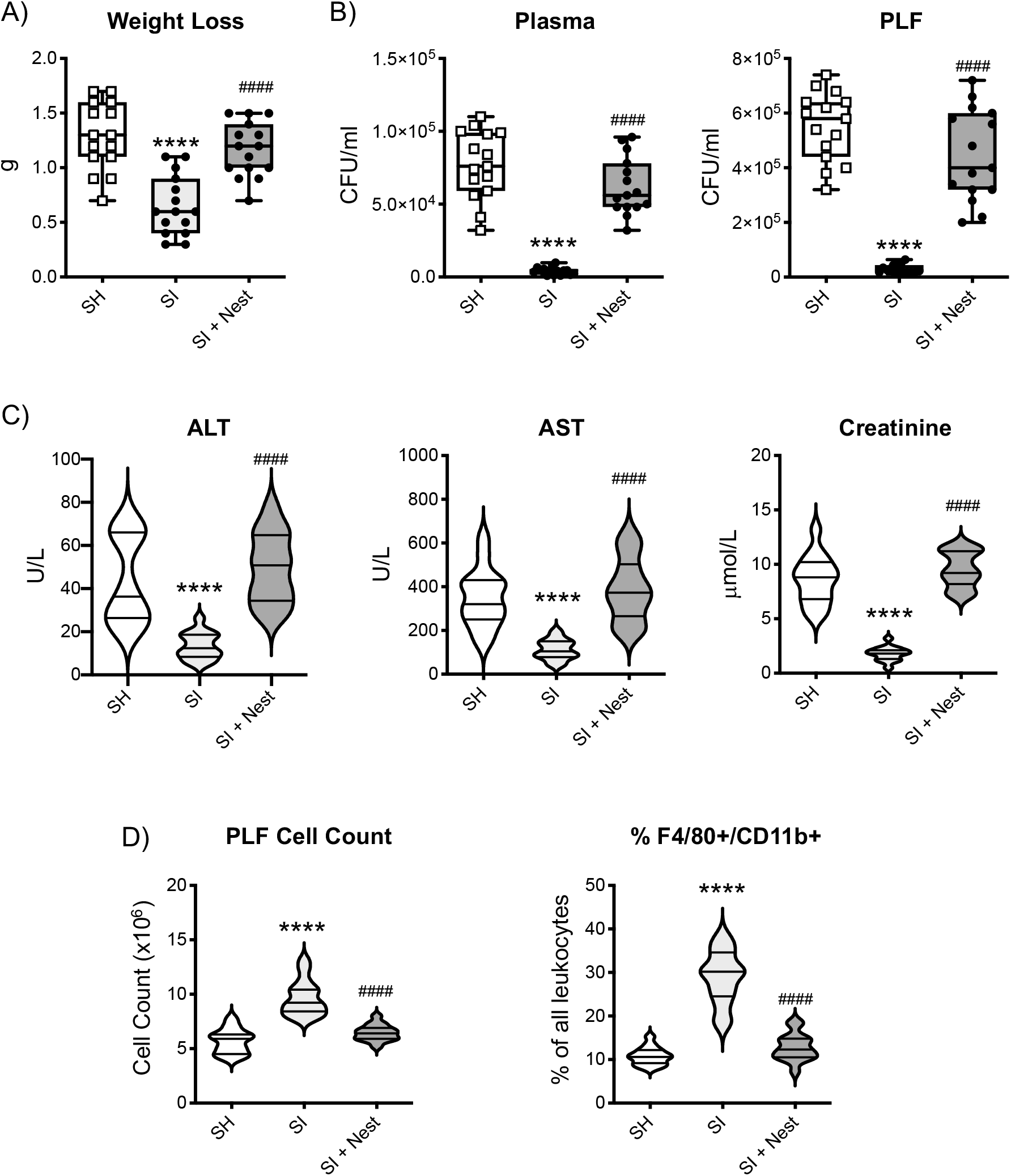
The effects of social isolation on bacterial clearance in CD-1 mice are reverted by the addition of an artificial nest. The box and whisker plots in (A) show the weight loss of socially isolated (SI), socially isolated + nest (SI+Nest) or socially housed (SH) CD-1 mice after 6 hrs from the challenged with 1×10^7^ CFU of *E*.*coli 06:K2:H1*. The box and whisker plots in (B) show the bacterial load of blood and peritoneal lavage fluids (PLF) of the same mice while the violin plots in (C) show the blood levels of ALT, AST and creatinine. The violin plots in (D) show the total number of peritoneal cells and their % of gated CD-11b/F4/80^high^ cells from the same mice. Each plot shows the median and the quartile of n=15 mice. Data are representative of n=3 independent experiments with similar results. ****p<0.0001 (One-way ANOVA) indicates significant values of SI mice compared to SH mice while ^####^p<0.0001 indicates significant values of SI+Nest mice compared to socially isolated.

## Discussion

The main goal of this study was to explore the impact of sole-living on the host immune response of CD-1 mice. Initially, we assessed if the increased state of anxiety-like behaviour shown by socially isolated mice would have any effect on the overall blood biochemistry and immune cell repertoire. Our results showed no significant changes in total leukocyte number or leukocyte subsets (lymphocytes, monocytes or neutrophils). The results of the blood biochemistry analysis provided the same results with both AST, ALT and creatinine being similar in SI and SH mice. Blood glucose levels were about 67% higher in SI mice - possibly suggesting increased gluconeogenic activity of the liver because of an increase in energy expenditure. Consistent with this hypothesis, a study on free-living African striped mice (Rhabdomys pumilio) has shown that solitary individuals have high levels of blood glucose compared to their social counterparts because of their higher energy expenditure (71).

As there was no overall significant change in the immune system cellularity of SI mice, we next sought to investigate if sole-living conditions would impact the way immune cells respond to pathogens, and to do so we challenged mice with LPS or live *E*.*Coli*. We used LPS to evaluate chiefly TLR-4-mediated inflammatory response and *E. Coli* to explore the ability of the host to clear an infection caused by live replicating bacteria. In the context of the response to LPS, our results concur with several seminal investigations (14, 33, 72-93) carried out in humans or primates by pioneers in field of research on social isolation like Cole, Cacioppo, Capitanio, Steptoe or Kiecolt-Glaser. In these studies, subjects with high levels of perceived loneliness or loneliness associated with a wide range of social or medical conditions showed an elevated inflammatory response or a significant upregulation of NFκB-responsive genes including IL-6 and TNF-α in their PBMC (29). Similarly, studies in primates subjected to social isolation because of low social status ranking showed a correlation between the heightened inflammatory response and their social status (94). In both humans and primates, this increased state of inflammation was linked to a skewing of TLR-4 downstream signalling towards the MyD88 antibacterial arm over the antiviral TRIF counterpart (79, 94).

We expanded on these studies through tests with live *E. Coli* that are not possible in humans. The intraperitoneal injection of *E. Coli* in mice is a useful experimental system to link bacteriemia with the expansion of the inflammatory response from local to systemic through the different stages of endotoxemia (95) i.e. Systemic Inflammatory Response Syndrome (SIRS), Septic Shock or Severe Sepsis, and finally Multiple Organ Dysfunction Syndrome (MODS) and death. Systemic (blood) and local (PLF) assessment of the bacteria load at the 6-hour time point demonstrated that SI mice were better placed in containing the bacteraemia and the subsequent systemic sepsis and organ damage. These results suggest that the functional outcome of the increased inflammatory response brought about by social isolation is to boost and accelerate bacterial clearance. Consistent with this hypothesis, we also found an increased number of mature phagocytic F4/80^+^/CD11b^+^ macrophages in the peritoneal cavity of SI mice. We think this is the reflection of the increased levels of inflammatory cytokines and chemokines found in these mice at earlier times (96, 97). In line with our data, a previous study on wound healing and infections has also shown that socially isolated mice had fewer bacteria in their wounds compared to SH mice. In contrast to our study, the authors reported that this was associated with a reduced expression of IL-1β and MCP-1 (98), possibly because they made their measurements at later time points rather than at their peak.

Microarray analysis of the whole blood of enriched mice revealed the modulation of a discreet set of genes (13 in total; 8 upregulated and 5 downregulated) that are known to foster an effective host-defensive inflammatory response (49). We wondered if these same genes would also be modulated by social isolation. More specifically, we hypothesised the existence of a selected group of immunomodulatory genes that would respond to changes in external living conditions and hence, be switched on and off by enriched or isolated housing. The results obtained provided a completely different answer and showed that sole living modulated the expression of genes that had very little resemblance or overlap in functions with that of the enriched environment. A large proportion of the DEG identified after social isolation were found to promote apoptosis or have proinflammatory roles. *CD55* was significantly higher in SI mice and has been reported to inhibit the activation of the complement pathway by affecting the C3 and C5 convertases (99-101). *CD55* is suppressed on neutrophils by NOD2-mediated signals during polymicrobial sepsis thus promoting C5a production which has been implicated in multiple organ failure, and cardiomyopathy during sepsis (102). *Xaf1* is a tumour suppressor gene involved in KIF1Bβ-mediated apoptosis where it acts as a molecular switch for p53 by promoting apoptosis as opposed to cell cycle arrest (103, 104). *Xaf1* gene expression decreased in blood at the onset of sepsis and recovered to normal levels within 48 hours in patients with low mortality scores but recovery of Xaf1 is delayed in those with high mortality scores (105). *Bach2*, a transcription factor, is involved in the promotion of T cell effector memory as well as B cell antibody class switching (106, 107). *Bach2* is highly expressed in functionally resting quiescent Treg cells and is down-regulated in activated T_reg_ cells during inflammation (108).

Zooming out from assessing the effects of sole-living at cellular and molecular levels, we were intrigued by the increased food intake of SI mice. Research on hyperphagia in the context of weight maintenance in mice suggests that this behaviour is likely to be secondary to increased energy expenditure to homeostatically control body temperature (109-111). This was rather interesting in light of a large body of recent work that links the immune response to metabolism and temperature control (112-114). Indeed, studies have shown that thermoneutral temperatures for mice (30-33°C) - as opposed to standard research laboratory temperatures of 22°C (a temperature that is tolerable for human caregivers) - simultaneously alter the host immune system and its metabolism (31– 34). A very interesting perspective by Christopher Karp in 2012 (115) highlighted how cold was deemed responsible for the allegedly increased resistance to LPS exhibited by experimental mice because of its ability to decrease myeloid cell differentiation.

This and many other considerations made us wonder: How do wild mice respond to changes to external temperature and specifically lower temperature? In mice (70, 116-119) and other species (120-127), huddling with other individuals is an effective way to reduce heat loss; it has indeed been considered “*a public good with a private cost*” (128). Behavioural studies have indeed shown that huddling in mice increases as the temperature lowers (117, 129). In neonatal rodents huddling reduces the cost of physiological thermoregulation and this behaviour is kept in adulthood despite the animal’s capacity to sustain a basal metabolism in isolation from the huddle (130). LPS or *E Coli*-induced sepsis in laboratory animals kept at ‘human-compatible temperature’ fail to develop fever and experience a hypothermic response (131) (see also **Video 1**) that makes them huddle to increase their body temperature. In our study, the provision of an artificial nest to SI mice offered a surrogate option to social thermoregulation. We chose this system instead of housing conditions with different (once again) ‘human-controlled’ temperatures as we wanted to give mice the freedom to adjust their temperature according to their needs. Recent studies have indeed challenged our understanding of a mouse’s thermoneutral zone (132-134) as their body temperature changes continuously and dynamically. Constant mouse body temperature monitoring has led to the discovery of thermoneutral points that changes according to the active or resting daily phases of mice (135). Last but not least, studies based on the use of a fixed ‘human-set’ thermoneutral housing condition have often reported contrasting effects on a wide range of disease conditions (113). As an example, from the perspective of an immune response to infection, thermoneutrality has been reported to both alleviate or exacerbate signs of disease (131, 136, 137).

This study is far from being exhaustive and opens several questions that would need to be addressed in the future. First and foremost, we wondered if this ‘primed’ state of the immune system brought about by social isolation would be indeed beneficial on the long term. A very large body of evidence suggests that a heightened basal level of immune activation might be responsible for the development of neurodegenerative diseases, psychiatric conditions and autoimmune disorders (138-141). This basal level of undetectable inflammation – also known as meta-inflammation-has been described to be most prevalent in subjects experiencing loneliness and social isolation (92, 142, 143). Secondly, we have not fully explored the link between immune cells, fat and heat regulation. For instance, recent studies have suggested that γ*/*δ T cells reside in fat tissues and regulate the response of mice to cold (144). We have noticed visible differences in the amount of visceral fat of mice housed in the 3 different conditions (SH, SI and SI+Nest; data not shown) but have not processed these tissues for detailed histological or cytofluorimetric analyses. We have also not performed a quantitative analysis of the brown adipose tissue (BAT) and white adipose tissue (WAT) as a large body of the literature has linked the inflammatory and the thermogenic response to the relative presence of these tissues (145). Finally, we used outbred CD-1 mice rather than widely used inbred C57BL/6 mice. We have consciously made this decision as it is now widely accepted that “*the use of inbred mice to model human disease is tantamount to using multiple copies of one individual*” (146, 147).

With all these considerations in mind, the results of this study still support an essential point that is embedded in our original research hypothesis: the immune system mirrors our lifestyle and what we are experiencing in life (10). Within this framework of reference, it is interesting to note that social thermoregulation in primates has been reported to be an evolutionary response to enhanced fitness i.e. enhanced survival and reproduction (65). At the same time, studies have suggested that primate fitness is positively correlated to total white blood cells and promiscuity (148, 149) thus supporting the idea of a continuum and a homeostatic balance between living conditions, behaviour and the immune system. The existence of this continuum would be of great scientific interest from a public health perspective. Especially in recent times when new approaches like social prescribing (150) and social determinants of health (151, 152) are been tested for their effectiveness in the prevention and treatment of a wide range of human conditions (153). We are indeed tempted to speculate that – like in the case of huddling in animals – there is a specific immunological and physiological response that is activated when social interactions and a sense of belonging are administered as a treatment. The availability of these results would provide a scientific base to the clinical effectiveness of these treatments while furthering appreciating the translational value and potential of the plasticity of the immune system.

## METHODS

### Animal Husbandry

Male CD-1 mice were purchased from Charles River and used for all experiments at 5 weeks of age. All animals were housed under standard conditions in individually ventilated enclosures with *ad libitum* access to food and water with a 12-hour light/dark cycle. All experiments undertaken were approved and performed according to the guidelines of the Ethical Committee for the Use of Animals, Bart’s and The London School of Medicine and Dentistry and the Home Office Regulations Act 1986, PPL 80/8714.

### Social Isolation Model

At 5 weeks of age, male CD-1 mice were assigned to social isolation or the control group social housing for 2 weeks. Mice assigned to social isolation were individually housed whilst those mice assigned to social housing had 5 mice that had been weaned together in them. All mice were placed into standard sized cages (**W × D × H:** 193 × 419 × 179 mm, Allentown) containing wood chip bedding and 2x strips of nesting material. Cages were cleaned 1 time per week and mice were only handled by one researcher. Mice were weighed at baseline and every week during cage cleaning. Food and water consumption were measured every 2 days.

### Behavioural Tests: Open Field and Light/Dark Shuttle Box

If not otherwise stated, tests were performed double-blinded during the light phase of the light-dark cycle, as previously described and recommended (154). All efforts were made to minimize mouse discomfort in these behavioural experiments. Mice were brought to the testing room at least 30 min before the start of the test session to allow habituation to the testing environment. Unless otherwise specified, standard lighting (about 50 lux) and quiet conditions were maintained throughout each experiment. Mice used for behavioural testing were not subjected to any other test or procedure. The open-field test is an ethologically based paradigm that provides objective measures of exploratory behaviour as well as a valid initial screen for anxiety-related behaviour in rodents and was carried out as previously described (155, 156). The apparatus consisted of a white PVC arena (50 cm × 30 cm × 20 cm) divided into 10 cm × 10 cm squares (n=15). The three central squares defined the “centre” region. Each mouse was placed in a corner square, facing the wall, and observed and recorded for 3 min. The total number of squares crossed (all four paws in), the total number of rears (defined as both front paws off the ground, but not as a part of grooming), and the number of centre crossings were recorded. The walls and floor of the arena were thoroughly cleaned between each trial.

The Light/Dark shuttle box test assesses the exploratory activity of mice and reflects the combination of hazard and risk avoidance (52) The apparatus consisted of a 45cm × 20 cm × 21 cm box, divided into two distinct compartments: one third (15 cm long) painted black, with a black lid on top, the remaining two-thirds painted white and uncovered. A 2.5 cm × 2.5 cm opening joined the two compartments. One side of the bright box was transparent to enable the behavioural assessment and the averseness of this compartment was increased by additional illumination supplied by a 50 W lamp placed 45 cm above the centre of the box floor. The test was performed in accordance with a previously published protocol (52). Each mouse was placed in the bright compartment, facing away from the opening and allowed to explore the box for 5 minutes. Dependent variables included the time spent in the light area, latency to cross to the dark area (all four paws in) and the total number of transitions between compartments. The apparatus was cleaned after each trial.

### Whole Blood Cellularity and Cytokine Profiling

Whole blood was aliquoted and send to Medical Research Council (MRC) Harwell for full blood count analysis on the Advia 2120 haematology analyser. Using a second aliquot of blood, plasma was generated by centrifugation of samples at 10,000 g for 5 minutes. Peritoneal lavage fluid (PLF) was spun down at 264 g for 5 minutes and the supernatant was removed. In the LPS model, plasma (tumour necrosis factor-α [TNF-α], interleukin-6 [IL-6], monocyte chemoattractant protein-1 [MCP-1], and interferon-γ [IFN-γ]) and PLF supernatant (TNF-α, IL-6, MCP-1, and KC) was sent to Labospace Ltd, Milan for cytokine analysis using multiplex assays. For all further models, TNF-α and IL-6 were measured according to the manufacturer’s instructions in plasma and peritoneal lavage supernatant using Novex ELISA Kits (Thermofisher Scientific Ltd.).

### Blood Biochemistry

Biochemical parameters (Aspartate transaminase [AST], alanine aminotransferase [ALT], creatinine and glucose) were measured in plasma at MRC Harwell on a Beckman Coulter AU680 clinical chemistry analyser. Corticosterone concentration in plasma was taken from mice after 2 weeks of social isolation was determined by enzyme-linked immunosorbent assay (ELISA) (Enzo) according to the manufacturer’s instructions.

### Microarray Analysis

Total RNA was extracted from heparinised blood of SI and SH mice (n=3 per housing group) using the RNeasy Protect Animal Blood Kit (Qiagen). Extracted RNA was hybridised at UCL genomics following standard Affymetrix protocols, using GeneChip Fluidics Station 450, and scanned using the Affymetrix GeneChip Scanner (Affymetrix, Santa Clara, CA, USA). Computational analysis was performed using Mac OS 10.6.8, and R version 3.1.0. Affymetrix platform MoGene_1.0st transcript cluster. Heatmap was generated by the function heatmap.2 of the CRAN package, gplots, using the complete-linkage clustering using the Euclidean distance. Data were normalized by robust multiarray average (RMA) of the Bioconductor package, affy. Relevant genes were filtered by excluding those without an Entrez ID and those with low expression levels less than 100 by non-logged value. T-statistics were applied across the data set using the Bioconductor package Limma considering the false discovery rate and differentially expressed genes were identified by p<0.05 (non-adjusted P-value).

### Real-time Polymerase Chain Reaction

Total RNA was extracted from blood (n=6 for each housing group) using the RNeasy Protect Animal Blood Kit (Qiagen) according to the manufacturer’s protocol. Reverse transcription of the extracted RNA was performed using 1μl of 0.5μg/μl OLIGO dT primers (Promega) and 1μl of RNase free-water and incubating samples at 70°C for 10 minutes and cooling for 5 minutes. 5x AMV reverse transcriptase buffer, RNAsinPlus, RNase free water, 10mM dnTP mix and AMV reverse transcriptase (all Promega) was added, and samples were incubated at 42°C for 60 minutes and 10 minutes at 70°C. Samples were loaded in triplicate and were run according to the manufacturer’s settings on a BioRad CFX Connect RT-PCR machine using power SYBR Green master mix (Thermo Fisher Scientific) and QuantiTech primers (Qiagen). RT-PCR. Data were analysed using the ΔΔC_T_ method with gene expression being normalised to Glyceraldehyde 3-phosphate dehydrogenase (GAPDH).

### Lipopolysaccharide (LPS) Induced inflammation

After 2 weeks of social housing or social isolation, mice were injected intraperitoneally with 15mg/kg of LPS (Sigma) resuspended in PBS. After 4 hours, mice were anaesthetized by isofluorane inhalation and blood was collected by cardiac puncture (anticoagulant: 3.2% sodium citrate). Animals were immediately sacrificed by CO_2_ asphyxiation and peritoneal lavage was performed using 2ml of 3mM EDTA-PBS. Cytokines in plasma (TNF-α, IL-6, MCP-1 and IFN-γ) and PLF supernatant (TNF-α, IL-6, MCP-1 and KC) was sent to Labospace Ltd, Milan for analysis using multiplex assays.

### *E*.*coli* Induced Sepsis

After 2 weeks of social housing or social isolation, mice were weighed before the induction of sepsis. Sepsis was induced *via* intraperitoneal (i.p.) injection of 1×10^7^ CFU of *E*.*coli 06:K2:H1*. At 6 hours, mice were weighed once again to allow sepsis-induced weight loss to be calculated. Mice were anaesthetized with isoflurane and cardiac puncture (anticoagulant: 3.2% sodium citrate) was performed before immediate sacrifice by CO_2_ asphyxiation. Peritoneal lavage was carried out using 2ml of PBS containing 3mM EDTA (PBS/EDTA). IL-6 and TNF-α were measured by ELISA and biochemical parameters (AST, ALT, creatinine and glucose) were measured as described previously.

Blood and peritoneal lavage fluid were diluted 1:1000 in sterile PBS and 50*μ*l of each was plated on LB agar plates. Plates were incubated overnight at 37°C and individual bacterial colonies were counted. CFU/ml was calculated by multiplying the number of colonies by the dilution factor and then dividing by the volume of the plate.

Cell numbers in the peritoneal lavage fluid were quantified *via a* haemocytometer. Samples were centrifuged for 5 minutes at 264 g and the pelleted leukocyte cells were resuspended in 200*μ*l of PBS/EDTA. Cells were washed in FACS buffer (PBS containing 5% fetal calf serum and 0.02% of NaN_2_) and blocked with 50*μ*l of CD-16/CD32 FcγIIR– blocking antibody (clone 93; eBioscience) in fluorescence-activated cell sorting (FACS) buffer. After 30 minutes, the block was washed off and the cells were stained with 50*μ*l of Anti-Mouse anti-CD-11b-BV785 (clone M1/70, Biolegend) and anti-F4/80-BV650 (clone BM8, Biolegend) in FACS buffer for 30 minutes at 4°C. Cells were then washed and fixed in 4% PFA and were acquired on an LSRFortessa flow cytometer (Becton Dickson). Analysis was done using FlowJo 7.0 software (Tree Star).

### Social Isolation with an Artificial Nest

CD-1 male mice were randomly assigned to social isolation, social isolation and an artificial nest or social housing for a period of 2 weeks. SI and SH cages were set up as previously described whilst mice assigned to be socially isolated with an artificial nest (SI + Nest) had cages containing 1x CD-1 mouse in standard size cage (W × D × H:193x 419x 179x mm, Allentown), 2x strips of nesting material, sawdust bedding up to 5cm in depth and 1x artificial nest. The latter was purchased from Amazon UK as “Blue Hammocks Hanging Bed House”, brand Hwydo, dimension H12cmxD12cmxW10cm seller.

## Statistics

According to the nature of the data obtained, a Student’s t-test (2-tailed), or a 1 or 2-way analysis of variance (ANOVA) was performed. Behavioural data were analysed via nonparametric analysis using the Mann-Whitney U test. All statistical analysis was performed using GraphPad PRISM software v8.0, with the exception of microarray analysis which was carried out as stated in 2.13.3 using the software package LIMMA (Bioconductor). Data were analysed for normality using the D’Agostino-Pearson omnibus normality test.

## Author Contributions

A.H helped with the designing of the studies, conducted experiments, acquired and analyzed data and contributed to writing the manuscript.

R.R & SB organized the preparation of the samples for the microarray analysis and contributed to some of the experiments.

M.O. performed the analysis of the microarray data.

D.C. & M.P provided advice on the experimental design and editing of the manuscript.

F.D. designed the study, conducted experiments, acquired and analyzed data and wrote the manuscript.

## ACKNOWLEDGMENTS

Alice Hamilton was funded by a British Heart Foundation PhD studentship.

We would like to thank Prof Jesmond Dalli (William Harvey Research Institute, Queen Mary University of London, UK) for providing the *E. Coli* used in the study and for the guidance on the sepsis protocol.

## Conflict of Interest statement

The authors have declared that no conflict of interest exists.

---

**Supplementary Figure 1.**
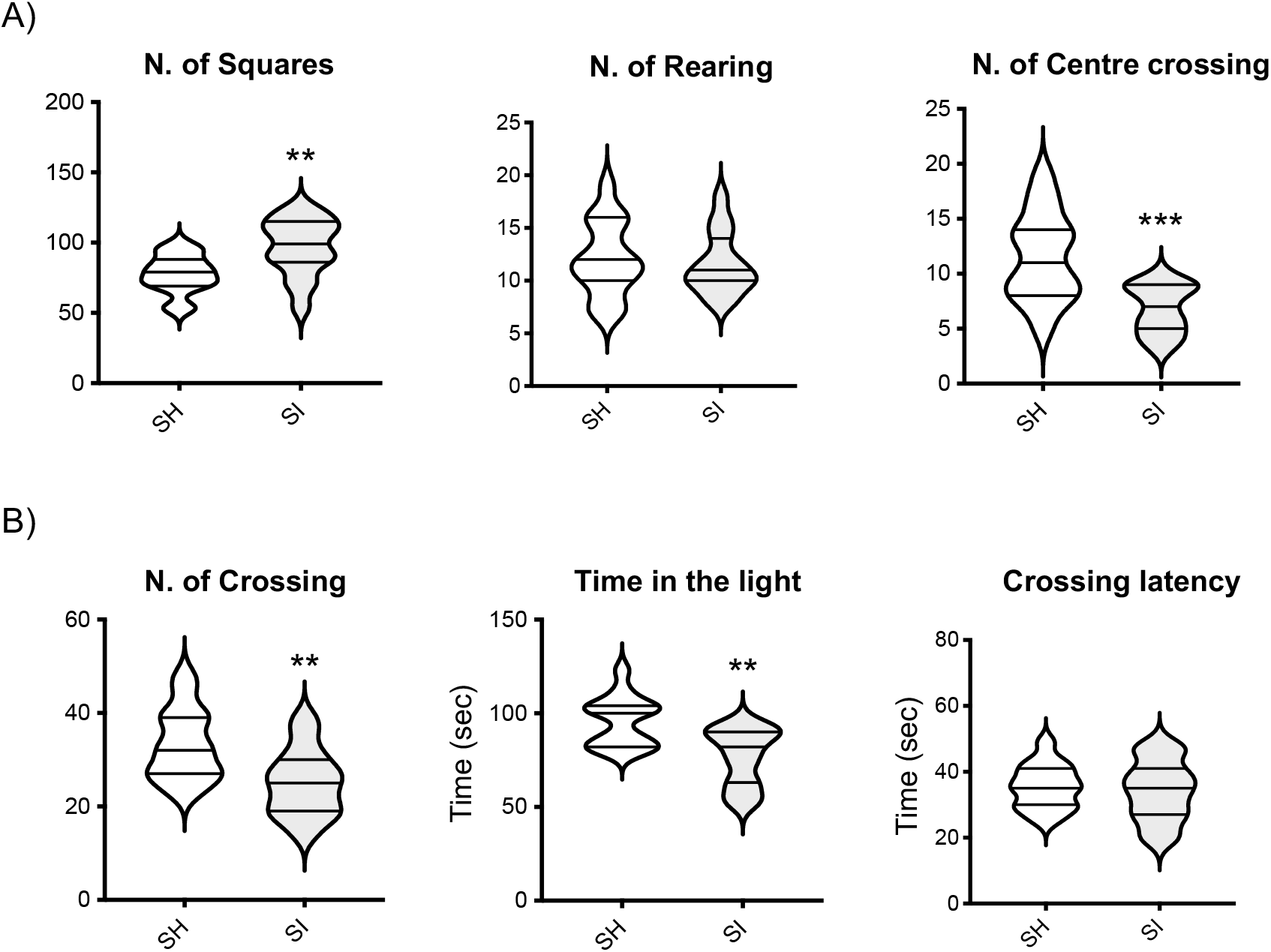
Effect of social isolation on the basal anxiety-like behaviour of CD-1 mice. The violin plots in (A) show the total number of squares crossed, the number of rears and the latency (seconds) to the first rear during a 5-minute session of CD-1 subjected to the open field test. The violin plots in (B) show the total time (seconds) spent in the lit area, latency (seconds) to first cross to the dark chamber and the total number of crossings during a 5-minute trial of CD-1 mice in the light/dark shuttle box test. Each plot shows the median and the quartile of n=15 mice. Data are representative of n=2 independent experiments with similar results. **p<0.01; ***p<0.001 indicate significant values compared to socially housed mice (Mann-Whitney U-test).

## Notes

### Competing Interest Statement

The authors have declared no competing interest.

